# New medium for the production of *Dermacentor reticulatus* and *Ixodes persulcatus* tick primary cell lines

**DOI:** 10.64898/2026.11.17.767464

**Authors:** Grigory G. Ananko, Alexey V. Kolosov

## Abstract

**Background:** The increasing prevalence of diseases caused by vector-borne infections is an important public health problem. Currently, over 1000 extant tick species have been identified worldwide, and approximately 10 % of them are known to be vectors of pathogens (protozoa, bacteria, and viruses), leading to diseases in animals and humans. Tick cell lines are increasingly being used in research on tick physiology and the pathogens of vector-borne infections.

To obtain and maintain tick cell lines, traditionally used L-15-based nutrient media. However, obtaining a continuous tick cell line remains a complex and time-consuming process. Therefore, it is important to develop new methods that can accelerate the process of obtaining tick cell cultures.

**Methods:** To establish primary tick cell lines, we used, for the first time, a ready-made DMEM/F12 + GlutaMAX culture medium supplemented with 20% fetal bovine serum.

**Results:** In this study, it was shown that DMEM/F12 + GlutaMAX nutrient medium is suitable for obtaining primary cell cultures of two species of ticks from two different genera of the family Ixodidae. Primary cell lines of the tick *Ixodes persulcatus* were obtained for the first time. We also managed to accelerate the process of obtaining subcultures: the first subcultures were obtained within 1–2 months of tick tissue isolation.

**Conclusions:** This new method, allows us to overcome several existing technical difficulties. It is likely that the DMEM/F12 + Glutamax medium can also be used to obtain and maintain tick cell lines of other species. Thus, an alternative to the traditionally used L-15-based media has emerged, allowing researchers to select the medium most suitable for their tasks. The rapid production of subcultures creates opportunities to conduct certain studies directly on primary cultures, without waiting for their transformation into an “immortal” culture.

**Highlights:**

- DMEM/F12 medium is suitable for obtaining tick primary cell lines.
- The new medium lets you get the first subculture faster.
- The alternative to the traditionally used L-15-based media has emerged.

## Introduction

Many arthropod species have a significant impact on both human quality of life and economic performance. Blood-sucking ticks can transmit pathogens that cause numerous human and animal diseases (Jongejan and Uilenberg, 2004). In particular, the subject of our study, *Dermacentor reticulatus*, is a known vector of infectious agents such as *Babesia canis, Babesia caballi* and spotted fever rickettsiae (Földvári et al., 2016). *Ixodes persulcatus* ticks are vectors of Lyme disease, recurrent fever, babesiosis, and viral tick-borne encephalitis (Belova et al., 2023). To develop measures to prevent and combat the spread of infections, it is necessary to study the persistence and reproduction of vector-borne pathogens in ticks. However, research on ticks involves technical and ethical difficulties related to their feeding and reproduction, as ticks are obligate hemophages at all stages of development, from larvae to adults. At the same time, many research tasks can be addressed via tick cell cultures.

The first continuous ixodid tick cell lines were established in 1975 from developing adult *Rhipicephalus appendiculatus* tissue and two of the original three lines are still in use today (Bell-Sakyi et al., 2018). Tick cell lines have been used to isolate, propagate, and study tick-borne viruses and bacteria (Zhou et al., 2018; Kurtti et al., 2016). They have also been used in studies of tick biology (Sterba et al., 2014), genomics (Miller et al., 2018), the microbiome (Bell-Sakyi and Attoui, 2016), tick vaccines (Contreras et al., 2017), and a number of other applications. Tick cell lines are also increasingly used as a cheaper and more ethical alternative to live ticks (Pohl et al., 2014; Bell-Sakyi et al., 2018).

According to Hartley et al. (2025), the Tick Cell Biobank of the University of Liverpool stores 67 continuous tick cell lines. The L15B growth medium, developed in 1989 (Munderloh and Kurtti, 1989), and several other modifications of L-15-based media have been used to generate and maintain most of the known tick cell lines (Holman and Ronald, 1980; Kurtti et al., 1983; Bell-Sakyi et al., 2022; Hartley et al., 2025). However, establishing continuous tick cell lines remains a lengthy and complex process, with historically low success rates (Bell-Sakyi et al., 2018). For example, first subcultures of tick primary cultures including L-15B medium were carried out after 2 months (Lima-Duarte et al., 2022), 5 months (Hartley et al., 2025), 6-12 months (Munderloh et al., 1994), or 1-6 years (Bell-Sakyi et al., 2022). This means that active cell proliferation occurs only after a very long delay (Bell-Sakyi et al., 2022).

In this study, we attempted to accelerate the process of obtaining tick tissue subcultures via DMEM/F12 + GlutaMAX culture medium supplemented with 20% fetal serum. The cells were isolated from the embryonic tissues (eggs) of two tick species: four primary cell lines of *Dermacentor reticulatus* and four primary cell lines of *Ixodes persulcatus* were obtained.

## Materials and methods

### Collection and Preparation

Engorged female ticks were collected from mammalian hosts (voles) in the field (Yakimenko et al., 2013). In April 2026, *Dermacentor reticulatus* ticks were collected in the Kinelsky district of Samara Region (European Russia), and *Ixodes persulcatus* ticks were collected in the Ordynsky district of Novosibirsk Region (western Siberia). The collected female ticks were placed one at a time into 50 ml plastic test tubes and sealed with a cotton-gauze stopper. To maintain humidity, a wet cotton swab was previously placed at the bottom of the tube. The ticks were transported to the laboratory during the day.

### Nutrient Medium

We used the ready-to-use nutrient medium DMEM/F12 + GlutaMAX™ 1 (1X) (GIBCO), a 1:1 mixture of two well-known media, DMEM (Dulbecco’s modified Eagle’s medium) and F12 (Ham). Fetal bovine serum (up to 20%) and gentamicin (up to 50 μg/ml) were added to the prepared nutrient medium. We used noninactivated fetal bovine serum, FBS100SA (DIA-M).

### Egg preparation

The container with the tick female was placed in a thermostat and incubated under optimal conditions for oviposition (26 °C, 90–95% humidity). A 50 ml glass bottle, covered with a piece of cotton fabric reinforced with an elastic band, was used as a container. The ticks were examined daily for the commencement of oviposition and subsequent embryonic development. The eggs were incubated under the same conditions until sampling.

### Isolation of primary cells from embryonic tissues

To obtain one *D. reticulatus* primary cell culture, oviposition of one female was used: 14 days after the start of oviposition, egg sample weights ranged from 0.05 to 0.11 g. To obtain one *I. persulcatus* primary cell culture, oviposition of one female was used: seven days after the start of oviposition, egg sample weights ranged from 0.09 to 0.15 g.

The eggs were placed in a cell strainer (100-μm pore size) and sterilized by immersing the strainer with the eggs in a freshly prepared 3% hydrogen peroxide solution for 3 minutes. The eggs (in the strainer) were then washed successively in two portions of sterile distilled water. The strainer with the eggs was placed on the sterile surface of a Petri dish, 200 μl of the medium was added, and a homogenate was prepared: the eggs were crushed directly on the strainer with the flat end of a pestle. The homogenate was then washed through the strainer with the prepared culture medium via a pipette. Thus, we removed large tissue fragments and whole eggs that remained on the strainer. The resulting cell suspension, 5 ml in volume, was transferred to Cell and Tissue Culture Flasks (BIOFIL surface treated) with a surface area of 25 cm^2^. The culture flasks were tightly sealed.

### Cell line maintenance procedure

The sealed culture flasks were incubated at 28°C in ambient air. Cell multiplication was monitored by examining the cultures by inverted microscope. Thirty to 50% of the culture medium was replaced weekly: a portion of the old medium was carefully removed with a pipette (the cells remained at the bottom of the flask), and fresh culture medium was added. Subcultures, as the cell density significantly increased, were performed. The contents of the original flask were diluted 2 times, when the cell concentration is increased to 2 ×105 cells/mlr: some cells retained in the parent flask and some transferred to daughter flask. Further subcultures were maintained in 25 cm^2^. The primary cell line is a mixture of loosely attached and nonattached cells. Therefore, before sampling, the cells were removed from the plastic surface via a 240 mm long Cell scraper S (TPP, Switzerland). If necessary, the selected sample was treated with a trypsin solution to break down the cellular conglomerates. A 30 µl culture sample was transferred to a 96-well plate and topped with a 0.15% trypsin solution preheated to 30°C (sample to trypsin ratio 1:5). The process was monitored using an inverted microscope; trypsinization typically lasts 10-20 minutes. The cell concentration was calculated via a Goryaev chamber after pre-staining with trypan blue: three grids for each sample; in tables present the average value ± standard deviation.

## Results

*D. reticulatus* tick primary_cell lines were isolated from embryonic tissues. A total of 7 cell lines were isolated from eggs of 7 females, but 3 cell lines were lost due were lost to contamination. The resulting homogenate, which was passed through a sieve (100 µm pore size), contained both individual cells and clumps containing a few to several dozen cells (Fig. 1a). The resulting cell lines were primarily suspension cultures, meaning that they were easily removed from the plastic surface by shaking the flask. However, some cells temporarily adhered to the plastic surface and were removed with a silicone scraper.

**Fig 1.**
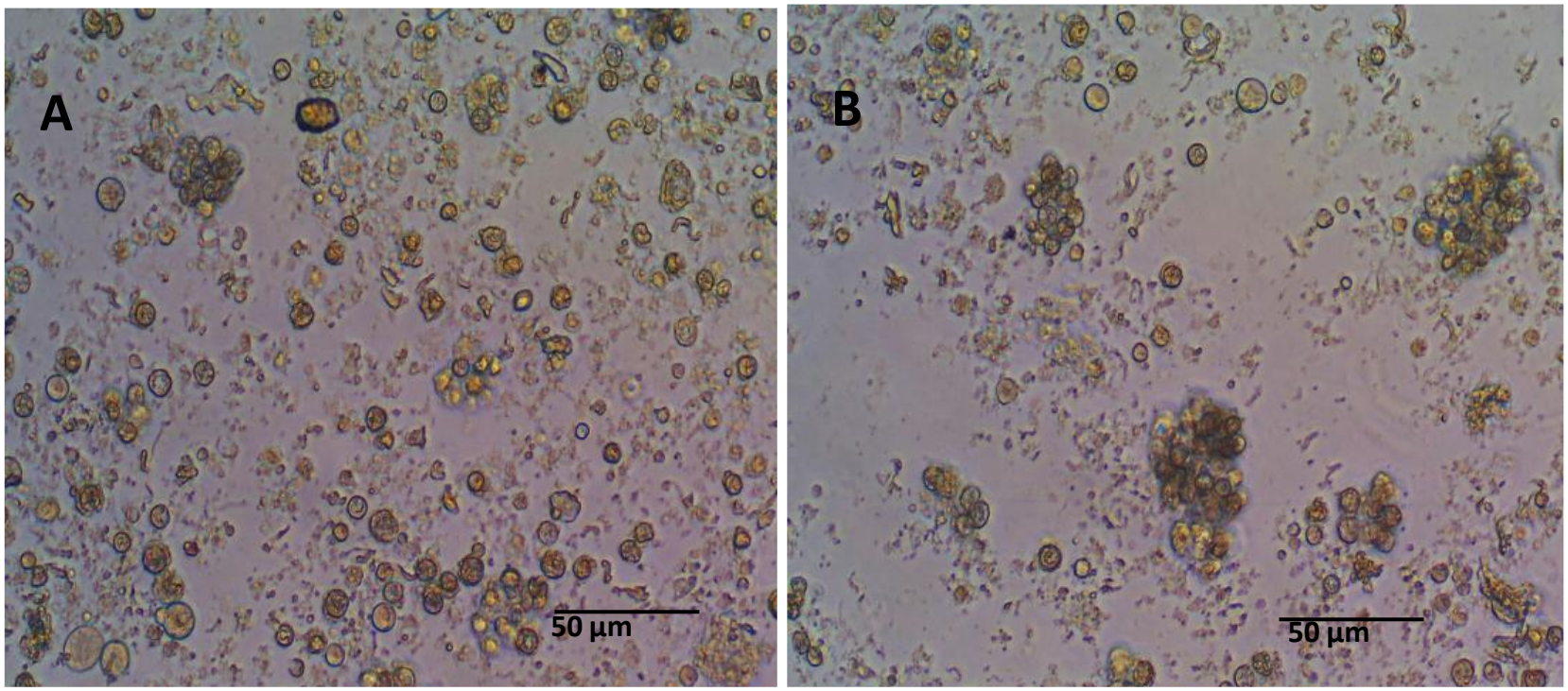
Primary tick cell lines: A - *Dermacentor reticulatus* (DR1); B - *Ixodes persulcatus* (IP1).

Among the primary *D. reticulatus* cell lines, the cell size varied from 5 to 45 μm. In the DR1 and DR2 primary cell lines, a significant increase in cell number was observed within 4 weeks of isolation (Fig. 2). The first subculture was carried out at 28 days in the DR2 line. Compared with the other lines, this line also yielded the greatest total number of cells (9.4×10^6^) over 12 weeks (Table 1). DR1, DR2 and DR3 cell lines were performed 3, 4 И 1 passages, respectively.

**Table 1.** Data on primary *D. reticulatus* cell lines.

| Cell line | Weight of egg sample, g | *Initial cell concentration, cell/ml ×10<sup>5</sup> | ** Initial number of cells ×10<sup>5</sup> | *** Final number of cells ×10<sup>5</sup> | **** Average doubling time, days |
|---|---|---|---|---|---|
| DR1 | 0.11 | 0.62±0.07 | 3.1 | 62 | 20.8 |
| DR2 | 0.11 | 0.54±0.05 | 2.7 | 94 | 17.5 |
| DR3 | 0.09 | 0.40±0.06 | 2.0 | 19 | 27.6 |
| DR4 | 0.05 | 0.23±0.05 | 1.1 | 2.5 | 75.7 |

\*Cell concentration on the day of isolation in a flask containing 5 ml of medium.
\*\*Total number of cells in 5 ml of medium on the day of isolation.
\*\*\*Final cell number from isolation: sum of all subcultures obtained over 12 weeks (84 days) in the specified cell line.
\*\*\*\*Average cell doubling time over the entire period (84 days).

**Fig 2.**
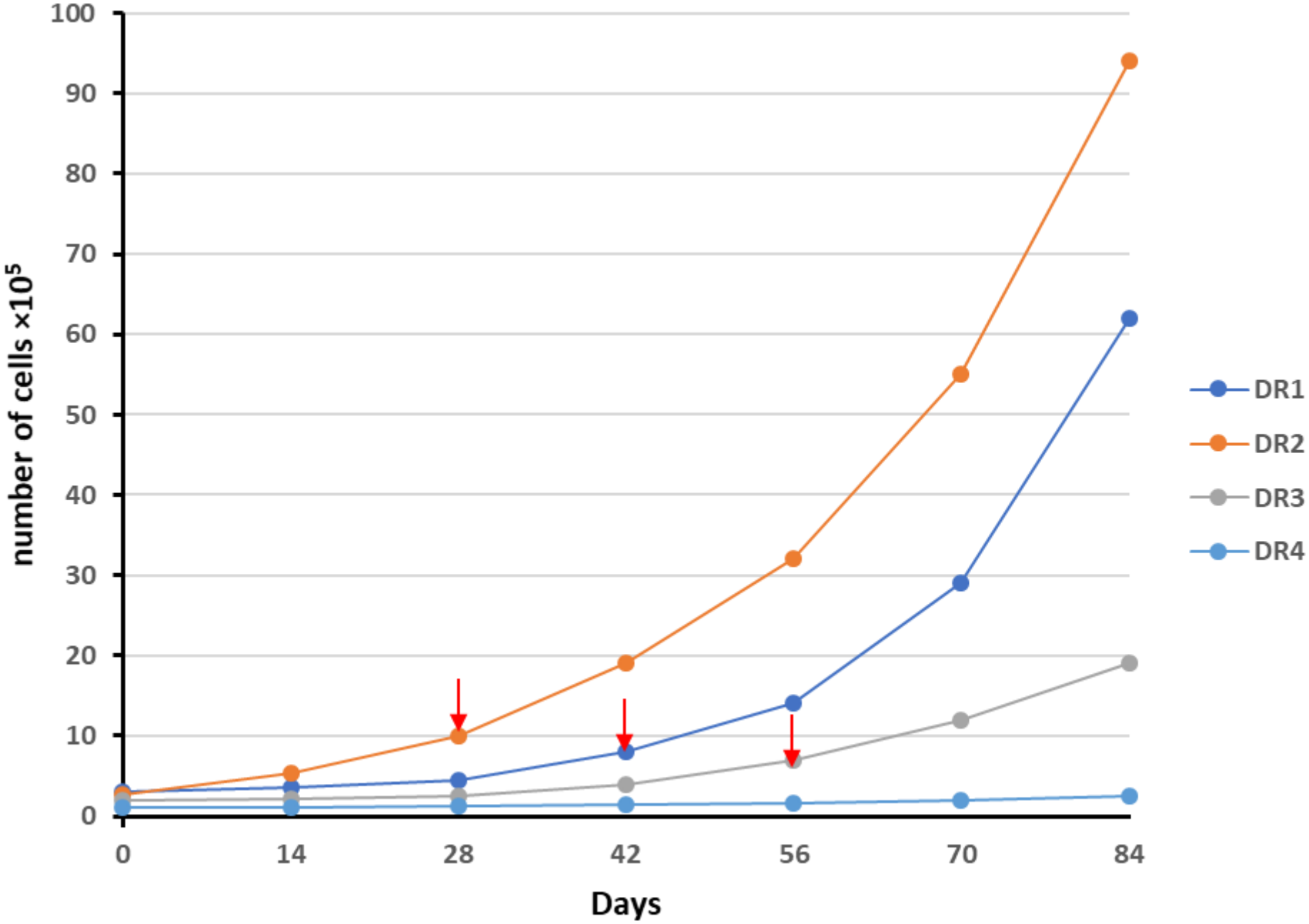
The dynamics of the proliferation of *Dermacentor reticulatus* primary cell lines (DR1, DR2, DR3, DR4) from the time of isolation from tick embryonic tissue. The cell number on the graph is the sum of the cell numbers across all subcultures obtained to date. The moment at which the first subculture is obtained is indicated by the red arrow.

*Ixodes persulcatus* tick primary cell lines were obtained for the first time: they were also isolated from embryonic tissues (Fig. 1b). A total of 5 cell lines were isolated from eggs of 5 females, but 1 cell line were lost due were lost to contamination. Unlike the method used to obtain *D. reticulatus* cell lines, we used eggs at an earlier stage of development, seven days after the start of oviposition. In addition, the average weight of the egg samples used for *I. persulcatus* cell isolation was greater, ranging from 0.09 to 0.15 g (Table 2). This is likely why the initial cell concentration in all *I. persulcatus* lines was relatively high. In the primary cell lines of *I. persulcatus*, the cell size varied from 5 to 30 μm. An increase in cell number was noticeable in all lines after just two weeks, especially in the IP1 line, where the first subculture was obtained after just three weeks (Fig. 3). IP1, IP2 and IP4 cell lines were performed 3, 3 И 1 passages, respectively.

**Table 2.** Data on primary *I. persulcatus* cell lines.

| Cell line | Weight of egg sample, g | *Initial cell concentration, cell/ml ×10<sup>5</sup> | ** Initial number of cells ×10<sup>5</sup> | *** Final number of cells ×10<sup>5</sup> | **** Average doubling time, days |
|---|---|---|---|---|---|
| IP1 | 0.15 | 0.78±0.10 | 3.9 | 36 | 28.0 |
| IP2 | 0.14 | 0.64±0.08 | 3.2 | 29 | 28.2 |
| IP3 | 0.10 | 0.52±0.06 | 2.6 | 8 | 55.3 |
| IP4 | 0.09 | 0.50±0.04 | 2.5 | 17 | 32.4 |

\*Cell concentration on the day of isolation in a flask containing 5 ml of medium.
\*\*Total number of cells in 5 ml of medium on the day of isolation.
\*\*\*Final cell number from isolation: sum of all subcultures obtained over 12 weeks (84 days) in the specified cell line.
\*\*\*\*Average cell doubling time over the entire period (84 days).

**Fig 3.**
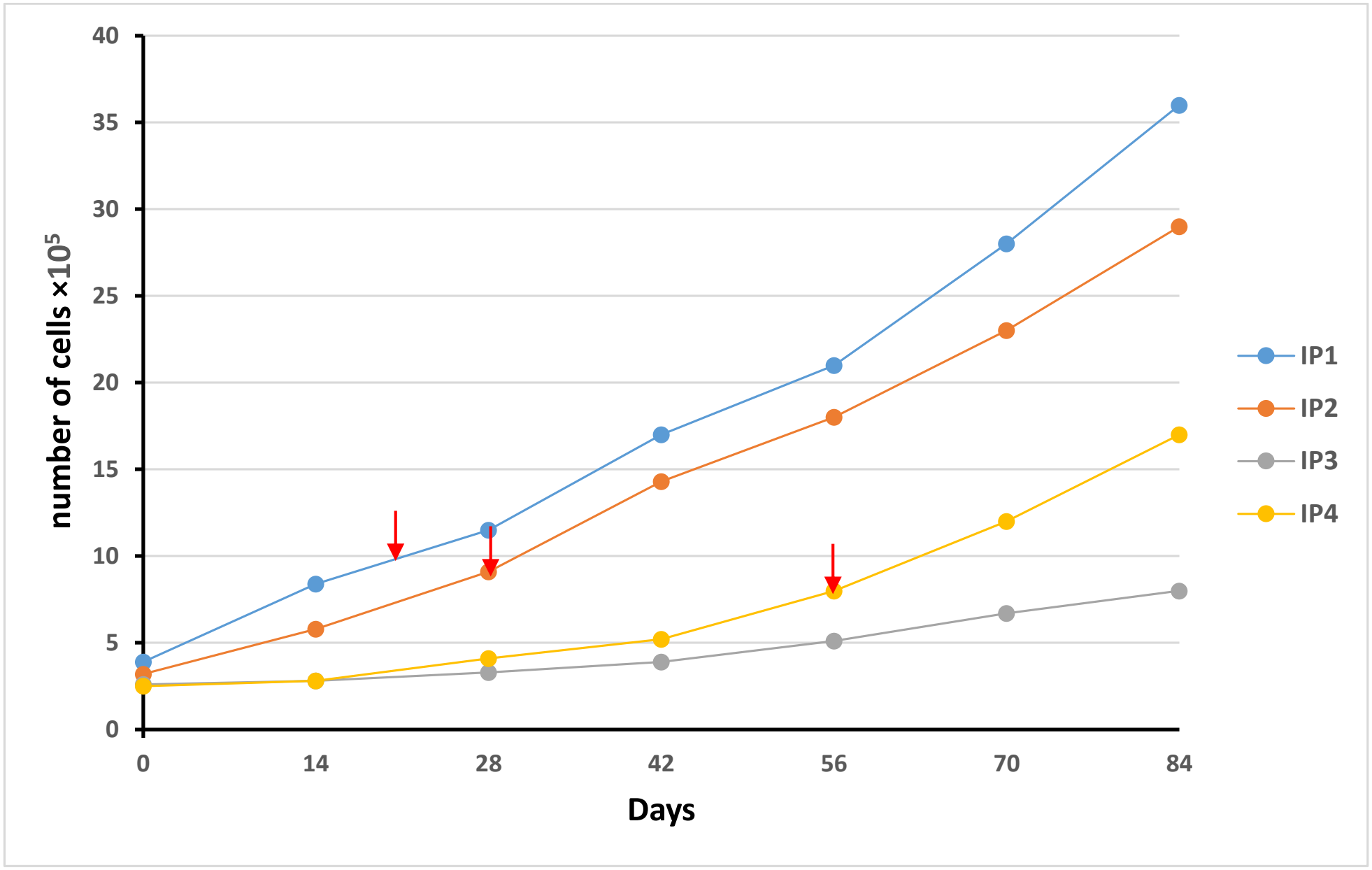
The dynamics of the proliferation of *Ixodes persulcatus* primary cell lines (IP1, IP2, IP3, IP4) from the time of isolation from tick embryonic tissue. The cell number on the graph is the sum of the cell numbers across all subcultures obtained to date. The moment at which the first subculture is obtained is indicated by the red arrow.

Comparing the doubling times of the best *D. reticulatus* (DR1 and DR2) and *I. persulcatus* (IP1 and IP2) lines, we see that the average cell doubling time of the first species (19.15±1.62) is almost 1.5 times shorter than the average cell doubling time of the second species (28.1±1.62).

## Discussion

L-15B medium, which is frequently used for the isolation and maintenance of tick cells, was created almost 40 years ago (Munderloh and Kurtti, 1989). It is based on Leibovitz L-15 medium (Leibovitz, 1963) supplemented with glucose, minerals, vitamins, tryptose-phosphate broth, cholesterol, amino acids, and tricarboxylic acids. Using this medium, *Ixodes scapularis* tick cell lines were first isolated (Munderloh et al., 1994). Subsequently, several modifications of the L-15 medium were created (H-Lac, L-15/MEM, and L-15B300), to which various concentrations of fetal bovine serum were added. Using these media, dozens of new tick cell lines have been generated (Bell-Sakyi et al., 2018; Bell-Sakyi et al., 2022).

However, the known procedure for producing new continuous tick cell lines remains complex and time-consuming, with a low success rate (Bell-Sakyi et al., 2022). To obtain a primary cell line, a homogenate of tick embryonic tissues (eggs) is typically used (Bell-Sakyi et al., 2018). The homogenate, consisting of tick tissue fragments, is then incubated in a nutrient fresh medium: a portion of the nutrient medium is replaced with fresh media weekly, and the primary culture is inspected by inverted microscope. The first subcultures of tick primary cultures including L-15B medium were carried out after 2 months (Lima-Duarte et al., 2022), 5 months (Hartley et al., 2025), or 1-6 years (Bell-Sakyi et al., 2022). At this stage, the risk of losing the primary line is high: when a primary culture is continuously maintained for several years, even with highly qualified researchers, it is not insured against accidental death.

We accelerated the process of obtaining subcultures by using a different nutrient medium, DMEM/F12 + GlutaMAX, with 20% fetal bovine serum, to isolate and maintain tick cell lines. It is a 1:1 mixture of two well-known media, DMEM and F12, with a number of additives. DMEM is a modification of Eagle’s minimal essential medium (MEM) and contains four times more amino acids and vitamins, as well as various additives that improve cell growth. DMEM, along with MEM, is widely used in mammalian cell culture. We used noninactivated fetal bovine serum. The question of which serum is better to use, heat-inactivated or not, has been little studied; previous studies on mammalian cells (Leshem et al., 1999; Bruinink et al., 2004) did not reveal any visible advantages of using heat-inactivated whey.

Using a ready-to-use culture medium (DMEM/F12 + GlutaMAX), we accelerated the production of the first subculture of primary *Dermacentor reticulatus* cell lines to 1-2 months. The cells likely began proliferating almost immediately after isolation. We also obtained similar results for primary *Ixodes persulcatus* cell lines: the first subcultures were obtained 3–4 weeks after isolation of tick tissue.

Perhaps there is a positive correlation between the initial cell concentration in the primary culture and the subsequent rate of cell proliferation. The higher the initial cell concentration (on the day of isolation), the earlier the increase in cell number was observed, and the greater the total number of cells obtained in each line over 84 days (Table 1). The initial concentration, in turn, positively correlates with the number of eggs (sample weight) from which the primary line was derived. For example, in the DR4 line, where the initial cell concentration was significantly lower (than in the DR1 line), 0.23 × 10^5^ cells/mL, the increase in cell number was very slow (Fig. 1), and no subcultures were obtained from this line during the first 84 days.

At the time of publication, the cell cultures were monitored for approximately 6 months. Although the cells proliferated actively after isolation, cell transformation apparently did not occur; that is, the resulting cell lines cannot yet be considered “continuous.” This conclusion is supported by the fact that between the 4th and 6th months of observation, all eight cell lines presented a significant reduction in the cell proliferation rate — by a factor of 2–3 (data not provided). A given cell line can be considered “continuous” as soon as a regular subculture rate has been achieved (Kurtti and Munderloh, 1982).

The use of DMEM/F12 medium simplifies the procedure for obtaining primary tick cell lines by using a ready-made nutrient medium: DMEM/F12 + GlutaMAX nutrient medium contains all the necessary components, except for serum and antibiotics (if necessary). In addition, owing to the active proliferation of cells almost immediately after tissue isolation, primary cells can be used for some studies where there is no need to use continuous cell line. In our experience, within 3 months of isolation, up to 10^7^ cells can be obtained from one line, which can be used for various research purposes. For example, for virological research. Overall, the new technique minimizes a number of existing technical difficulties (Wang and Cull, 2025) that limit the number of researchers interested in working with tick cells.

## Conclusions

In this study, we demonstrated that nutrient medium DMEM/F12 + GlutaMAX is suitable for producing primary cell cultures of two tick species belonging to two different genera of the family Ixodidae. We also succeeded in significantly accelerating the process of obtaining initial subcultures. It is likely that DMEM/F12 + GlutaMAX can also be used to obtain and maintain tick cell lines of other species. Thus, an alternative to the frequently used L-15-based medifa has emerged, allowing researchers to select the medium most suitable for their tasks. We hope that the use of this new medium will also help reduce the time required to obtain a continuous cell line.

## Author Contributions

**Grigory G. Ananko:** Conceptualization, Methodology, Investigation, Data curation, Formal analysis, Writing, Original draft preparation. **Aleksei V. Kolosov**: Investigation, Data curation, Writing. All authors have read and agreed to the published version of the manuscript.

## Ethics declarations

### Ethics approval and consent to participate

Not applicable.

### Consent for publication

Not applicable.

